# Data-driven identification of functional networks in artificial and biological neural networks

**DOI:** 10.1101/2025.11.18.688989

**Authors:** Karla Ivankovic, Anastasios Dimou, Justo Montoya-Gálvez, Riccardo Zucca, Manuel Valero, Alessandro Principe

**Affiliations:** Hospital del Mar Research Institute, 08003, Barcelona, Spain; Universitat Pompeu Fabra (UPF), 08003, Barcelona, Spain; Centre for Research and Technology Hellas (CERTH), GR 57001 Thermi, Thessaloniki, Greece

**Author notes:** **Corresponding author:** Alessandro Principe, Address: Hospital del Mar, Passeig Marítim 25-29, 08003 Barcelona, Spain.

**Keywords:** functional network, artificial neural network, EEG, prediction error, neural representations, parallel distributed processing

## Abstract

Understanding how the brain represents information is a central challenge in neuroscience and a practical bottleneck for brain-computer interfaces. Existing analytical tools cannot identify neural representations directly from neural activity data. We introduce MultiPEC, a data-driven method that discovers neural representations by quantifying how sets of signals jointly reduce context prediction error. Applied to artificial neural networks, MultiPEC uncovered class-specific subnetworks whose targeted ablation disproportionately impaired performance, demonstrating their causal role in feature recognition. Applied to EEG recordings from 24 participants, MultiPEC revealed functional signatures of auditory and visual processing. Classification analyses showed that MultiPEC captured fine-grained stimulus submodalities (levels of stimulus meaningfulness) more effectively than broad stimulus domains, highlighting the context-sensitive nature of neural representations. Together, these results establish MultiPEC as a scalable approach for identifying data driven (natural) representations in both biological and artificial systems, with potential applications in adaptive neurotechnology, clinical diagnostics and cognitive neuroscience.

**Highlights:**

- MultiPEC identifies neural representations by extracting information clusters from neural activity data.
- EEG analyses reveal individualized, condition-specific functional networks across auditory and visual modalities.
- In convolutional neural networks, MultiPEC uncovers class-specific networks critical for model performance.
- Provides a data-driven, hypothesis-free framework for mapping neural representations in biological and artificial systems.

## 1. Introduction

Understanding how the brain represents information about the world remains one of the central challenges in neuroscience (Mathis et al., 2024). Distributed neuronal activities are organized into coherent functional representations that support behavior and cognition Click or tap here to enter text.. However, it remains unclear how we can isolate these representations directly from neural activity data. This question is not only fundamental for basic neuroscience, but also for brain-computer interface (BCI) applications (Pandarinath et al., 2015; Sussillo et al., 2012), where decoding neural activity requires tracking internal representations as the brain itself adapts and reorganizes (Chang et al., 2024; Lebedev & Nicolelis, 2006; Sadtler et al., 2014).

Neural representations have been studied across scales. Optogenetic experiments demonstrated causal roles for neuronal ensembles in specific behaviors (Carrillo-Reid et al., 2019; Iurilli & Datta, 2017; Panzeri et al., 2022), and large-scale imaging and electrophysiology revealed distributed networks whose activity correlates with cognition (Combrisson et al., 2025; Farzmahdi et al., 2025). Mechanistic and statistical approaches have advanced our understanding of the mathematical principles underlying neural encoding, while also enabling practical applications in BCIs (Mathis et al., 2024). Practically all BCI approaches relate neural activity to explicit external variables (e.g., motor output) (Ding et al., 2025; Metzger et al., 2023). In contrast, emerging brain foundation models offer a data-driven alternative by uncovering structure directly from neural recordings, yielding latent representations such as manifolds, subspaces, and embeddings that reflect how neural populations organize their activity (Wang et al., 2025). However, these latent representations do not map directly onto recording channels, and no current approach automatically identifies subnetworks of electrodes or neurons that encode specific stimulus features.

Conventional analytic frameworks also present important limitations. Functional connectivity studies in humans are correlational, and traditionally look at pair-wise signal relationships, which misses the distributed nature of brain networks (Liu et al., 2025; Santoro et al., 2024; Sparacino et al., 2023). Dimensionality reduction methods such as principal component analysis (PCA) (Cunningham & Yu, 2014; Lever et al., 2017) can disassociate naturally coupled signals and obscure redundancies that may be part of a representation but do not explore multidimensional associations. Supervised decoding methods, such as multi-voxel pattern analysis, depend on predefined labels, limiting their capacity to uncover the full repertoire of representational structures across scales (Schreiber & Krekelberg, 2013).

A deeper understanding of brain function requires tools that can identify neural representations directly from neural activity, without a priori assumptions about stimulus categories or spatial scale. Taken together, three methodological challenges remain: (i) quantifying the information content of neural activity, (ii) identifying data models (representations) directly from neural recordings without predefined labels, and (iii) enabling empirical validation of discovered models beyond simple correlation.

Here we introduce MultiPEC (Multi-node Prediction Error Connectivity), a computational method for data-driven, systematic identification of neural representations. MultiPEC builds upon PEC (Principe et al., 2019), a metric that quantifies functional connectivity by measuring the prediction error of context modeled time series. Extending this principle to multiple nodes, MultiPEC systematically evaluates combinations of nodes and identifies those that jointly minimize prediction error. By monitoring when additional nodes no longer reduce the error, the method defines a boundary of a candidate functional module. Unlike correlation or synchrony measures, MultiPEC can capture predictive and asynchronous dependencies, and unlike PCA, it preserves redundancies that likely characterize distributed coding. The outputs are explicit node sets that are directly amenable to follow-up testing (e.g., ablation or targeted perturbation) and to use as structural priors for downstream decoders.

We evaluate MultiPEC on artificial neural network and EEG data. Our findings demonstrate that MultiPEC uncovers neural representations in a scalable, data-driven manner that respects the complexity and individuality of brain function. By bridging current gaps in neural data analysis, MultiPEC opens new opportunities for exploring the principles of distributed coding in the brain and for building adaptive BCIs.

## 2. Methods

### 2.1. MultiPEC algorithm

We designed the MultiPEC algorithm to identify neural representations (data models) by extracting groups of nodes (EEG channels or convolutional neural network filters) that exhibit low prediction error connectivity (PEC) (Principe et al., 2019). PEC is a connectivity measure based on the context modeling of a compression algorithm, that quantifies how well the activity of one node can be predicted from the activity patterns of other nodes. Adding informative nodes reduces the prediction error, whereas adding irrelevant nodes increases it. Following the original implementation (Principe et al., 2019), PEC was computed by (i) binarizing each timeseries to retain temporal structure, (ii) serializing the bitstreams of multiple nodes into a single sequence, and (iii) estimating the average weighted context model prediction error using a context length of 6 × N (where N is the number of nodes). This yields a scalar PEC value for any candidate node group.

The resulting prediction error is used to define functional networks, without pre-specifying information categories. Pairwise PEC values are first computed for all directed nodes pairs, and for each pair, the direction with lower error is retained. Because an exhaustive search becomes computationally unfeasible for large node sets (the number of possible combinations exponentially increases), we leveraged the observation that the PEC of a starting node pair is highly predictive of the final network PEC (Pearson’s coefficient = 0.99; Fig S2). This allowed us to restrict the search space by grouping node pairs according to their position within the PEC distribution. Specifically, node pairs were binned into seven ranges defined by standard deviations (σ) below or above the median PEC. These ranges reflect the negatively skewed PEC distribution and limit the number of candidate combinations explored by MultiPEC. Within each group, the algorithm iteratively constructs networks by greedily expanding node groups starting from the pairwise seeds. At each step, the candidate node set updates, and the average PEC error of the resulting subnetwork is tracked. The expansion stops when adding more nodes no longer improved (i.e., increased) the error. Each run produces a network and its associated error evolution. This process repeats across all groups, generating multiple networks per subject or model. The resulting subnetworks reflect channel groupings with varying degrees of internal efficiency, as measured by compressive predictability. All subnetworks were stored for further analysis and comparison across stimulus types or data modalities.

### 2.2. Simulated binary series for controlled validation

To validate the fundamental principles of MultiPEC, we first applied the method to a synthetic dataset of simulated binary series. This experiment provided a simplified environment where the ground-truth structure of information was precisely defined, allowing direct evaluation of MultiPEC’s ability to recover known representations. The dataset consisted of deterministic symbol sequences designed to emulate neural activations corresponding to distinct informational categories. Sequences of characters were generated for each experimental round using randomized combinations of uppercase letters, numerical digits, and mixed alphanumeric sequences. For each round, three sets of sequences were constructed: Set A (uppercase letters), Set B (digits), and Set C (mix of uppercase letters and digits in equal ratios), with each set containing six unique sequences of eight characters. The characters were sampled randomly, ensuring non-redundant content across sets. Each of these base sequences was then expanded into longer streams through a two-stage repetition process. First, every character in a sequence was repeated ten times to amplify its influence. The sets were then shuffled and repeated across ten full cycles, with a baseline symbol (‘@’ and ‘=’) inserted between each unit to segment the stream. This procedure yielded a structured input of 6 × 8 × 10 × 10 = 4800 characters per set, per round.

Characters (letters, digits, and punctuation) were then encoded using either 5-bit or 6-bit binary schemes derived from the 8-bit ASCII (American Standard Code for Information Interchange). ASCII encodes 128 characters, while we only considered 36 (24 upper case letters, 10 single digits, and 2 punctuations). Including the redundant bits would lower the control we have over the experiment, as we wanted to limit the overlap between the sets. Therefore, we reduced the code to 5 and 6 bits, in two separate experiments, by removing the first bits of ASCII. The experiment with 6 bits had a little overlap between sets, as the first bit was shared between some characters. Each bit of the binary vector was mapped to a separate node, effectively distributing the information spatially across a network. For two-network simulations, bits were assigned to 10 or 12 nodes (for 5 or 6 bits per network, respectively), split evenly between Network A and Network B. For three-network simulations, the bits were distributed across 15 or 18 nodes, split between Networks A, B, and C. The assignment of binary information to networks depended on character type. Each set (A, B, or C) was encoded so that their information was embedded in the corresponding network (e.g., Set A’s characters encoded in Network A), while the other networks received randomized binary noise. This structure allowed controlled overlap and independence between networks. Baseline characters were encoded identically across all networks to simulate shared non-informative input.

The nodes were analyzed using the MultiPEC method to infer network structure. The identified networks were subsequently compared against the known ground truth to assess accuracy. Identified networks were classified into three categories: perfect prediction, subset, and imperfect prediction, based on the overlap with the ground truth. Perfect prediction included networks that perfectly matched one of the predefined networks. Subset included networks with nodes contained entirely within a single predefined network but did not include the entire predefined network. Imperfect prediction included any network that spanned across multiple predefined networks.

### 2.3. Convolutional neural network model

#### 2.3.1. Model architecture and training

To test the data-driven network segregation method, we used a convolutional neural network (CNN), which is well-suited for visual tasks due to its architecture that mimics the hierarchical processing found in the visual cortex (Hubel & Wiesel, 1962). We implemented a small model for classifying the MNIST (Modified National Institute of Standards and Technology) image dataset of handwritten digits from 0 to 9, which makes for a total of 10 distinct input classes, as per (Koehler, 2020) using PyTorch (Ansel et al., 2024). The CNN model consisted of two convolutional layers (conv1 and conv2) and two fully connected layers (Fig 1). The convolutional layer dimensions are determined by the input depth dimension (1 for grey scale), the number of kernels (10 for conv1, and 20 for conv2 and kernel dimensions (5×5). Detailed architecture can be found in Supplementary material. Training data were divided into batches of 64 images. The negative log likelihood loss function was used to measure the predicted probability distribution. Stochastic gradient descent (SGD) optimization algorithm was used to update the model parameters iteratively through backpropagation to minimize the loss function prediction error (learning rate was 0.01 and momentum was 0.5). Test data were divided into batches of 1,000 images. The model achieved an accuracy of 98% after 10 training epochs (Fig S2). Accuracy for class ‘0’, ‘1’ and ‘4’ was 99%, class ‘2’, ‘3’, ‘5’ and ‘6’ was 98%, and class ‘7’, ‘8’ and ‘9’ were 97%.

**Figure 1.**
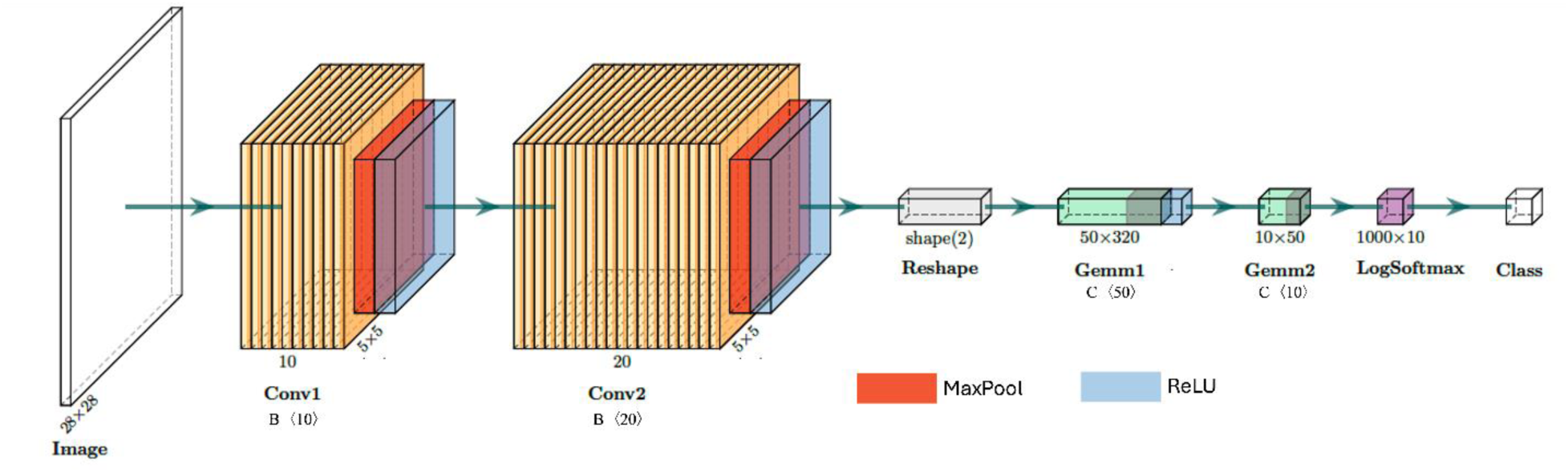
CNN model architecture. The model included two convolutional layers (Conv1 and Conv2) with ReLU activation (blue), pooling (red), dropout for regularization, and two fully connected layers (Gemm1 and Gemm2). Input were grayscale images of 28×28 pixels (MNIST). The convolutional layer dimensions are determined by the input depth dimension (1 for grey scale), the number of kernels (10 in the first, and 20 in the second layer) and kernel dimensions (5×5). The final output was a log SoftMax activation, providing the log-probabilities of each class.

#### 2.3.2. Activation preprocessing for MultiPEC analysis

To extract internal representations of the CNN model, we conducted a controlled experiment by continuing the training on a balanced subset of MNIST data. For each class, 20 images were selected from the training set, compiled into training batches with shuffling, in 5 training epochs. Dropout layers were deactivated during this phase to maintain deterministic outputs. At the end of each epoch, the model was evaluated on the MNIST test set to monitor performance. Feature maps were extracted for each epoch and batch. Each feature map was averaged spatially to produce a single scalar per filter per image. In this case, a kernel was considered a neural mass, comprised of neurons that share the learnable parameters. All filter-averaged activations were put in sequence to construct a 2D signal array (analogous to an EEG time series), with dimensions (number of filters, number of images), where number of filters was 30 and number of images 1,000. If null filters were detected, the process was halted to prevent propagation of invalid signals. Signals were binarized, and signals with all values equal to one were removed. We will refer to the averaged filters as “nodes”, in keeping with the neural network theme.

#### 2.3.3. Network evaluation via targeted pruning

To evaluate the functional role of identified networks, we applied targeted pruning of the CNN model and assessed the pruned model’s performance. For each network, a fresh instance of the trained model was loaded, and a custom binary mask was applied to zero out (i.e., prune) the weights associated with the specified indices of the network. The pruned model was then evaluated on the MNIST test set, yielding overall and per-class accuracy. The pruned model’s class accuracy was subtracted from the original model’s class accuracy to calculate the class accuracy change (Δ*A_c_*). The difference between maximum Δ*A_c_* and average Δ*A_c_* represented the network’s class specificity (*S*), with the set of all classes as *C* (Equation 1).

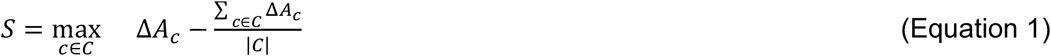

#### 2.3.4. Randomized pruning control

To evaluate whether specific groups of nodes show class-specific behavior, we conducted a randomized global pruning search, which consisted of 1,000 instances of pruning random groups of conv1 and conv2 nodes. Group size range was from four to seven nodes, matching the that of MultiPEC-identified networks. For each pruned model, test per-class accuracy was measured to compute class specificity (Equation 1).

### 2.4. EEG recordings

#### 2.4.1. Dataset overview and preprocessing

We analyzed a published EEG dataset by Orłowski and Bola, available on the Open Science Framework platform (https://osf.io/e93db/). The data set included 64-channel EEG recordings of neural responses to audiovisual stimuli with varying levels of meaningfulness, collected from 24 healthy adults. Participants were exposed to seven experimental conditions (tasks): original, scrambled, and noise versions of 30-second auditory (audiobook) and visual (cartoon) fragments, along with a resting-state condition. Stimuli manipulation preserves low-level sensory features while disrupting semantic content. Each condition was presented ten times in a randomized order. The study followed ethical guidelines and was approved by the Jagiellonian University ethics committee. The data were preprocessed as described in (Orłowski & Bola, 2023) kurtosis was used to identify artifact components after applying the independent component analysis. MultiPEC networks were processed for node pairs with PEC value lower than 3 × σ below the median.

### 2.4.2. Feature vectors

To interpret EEG networks in terms of underlying cognitive processes, each of the 62 EEG electrode channels (excluding references A1 and A2) was annotated with one or more cognitive functions based on its known neuroanatomical location and functional roles. These annotations were curated manually from literature, associating each channel with descriptions such as “motor planning”, “auditory processing”, “visual attention”, and others. A mapping was then constructed between standardized function labels and the set of channels that support them (Figure 2). For instance, network including channels ‘FC5’, ‘T7’, and ‘TP7’ would be annotated with auditory-motor integration, speech perception, auditory processing, language and speech and language integration. Channel-to-function mappings followed validated MRI-based projections of 10–20 electrodes onto cortical areas (Koessler et al., 2009).

**Figure 2.**
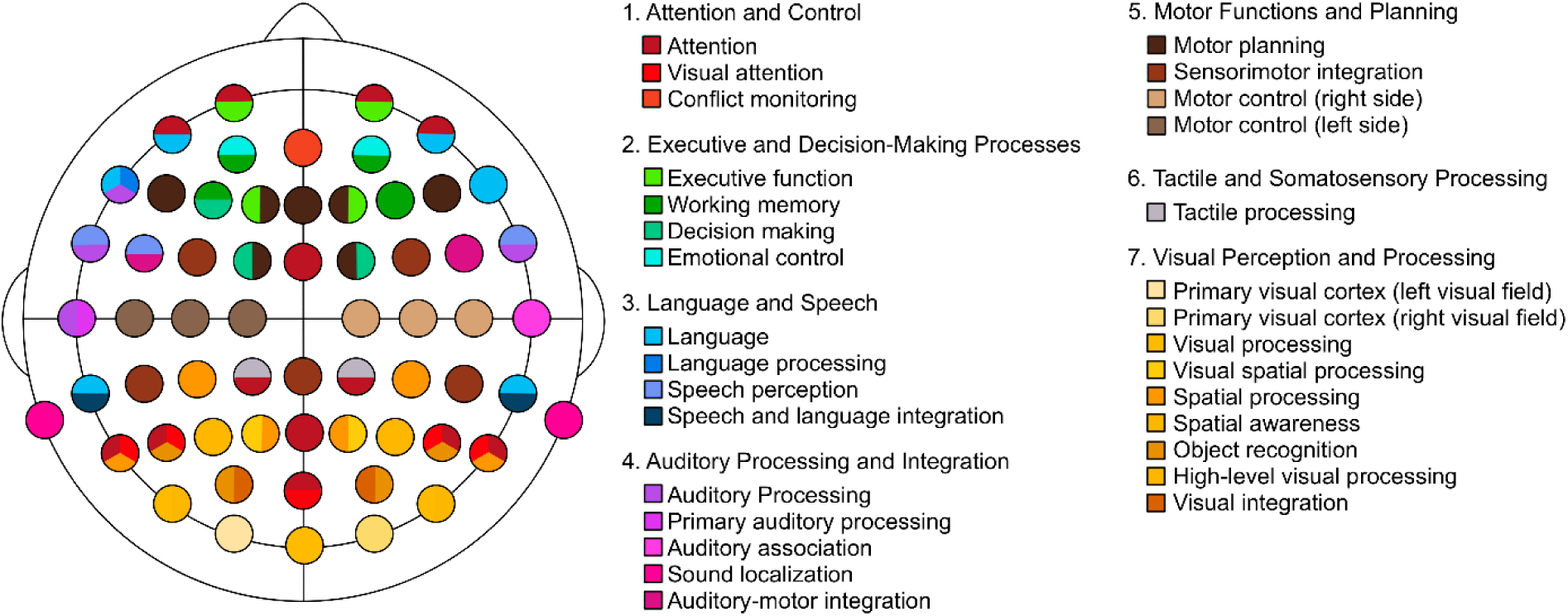
Mapping of cognitive functions to EEG channels for 64-channel scalp EEG.

Networks were transformed into PEC-weighted cognitive function vectors based on channel-level function mappings. First, for each network, a zero-initialized vector was created where each dimension represented a cognitive function from Figure 2. Network PEC value was normalized using reversed min-max normalization, such that lower PEC values received higher weights as per Equation 2.

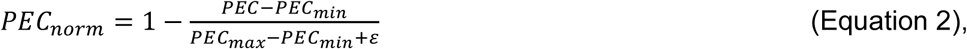

where ε =10^−6^ to avoid division by zero.

For every EEG channel in a network, we retrieved the associated cognitive functions and incremented the corresponding vector entries by the network’s normalized PEC. Each vector was indexed using a unique subject-task-network identifier. The resulting vectors were used for classification of tasks.

#### 2.4.3. Task classification

Classification was done for each subject separately. The feature matrix was first standardized using Z-score normalization. The linear support vector machine (SVM) classifier was used in a one-vs-rest configuration to accommodate the multi-class classification. The SVM was configured to address the substantial imbalance in the number of samples across classes (varying number of identified networks per task). Each class was assigned a weight that was inversely proportional to its frequency, increasing the influence of underrepresented classes and reducing that of overrepresented ones during learning. To minimize the bias towards most represented classes, stratified k-fold cross-validation was applied (k=5), which maintained consistent class distributions in both the training and testing folds. Only classes represented by at least five samples (equal to the number of folds) were retained. During each fold, the classifier was trained on the training subset and evaluated on the held-out fold.

In the first classification analysis, we aimed to emphasize the modality-based distinctions. The three audio-only conditions were grouped into a single “audio” class, while the three video-only conditions were grouped into a single “video” class. The resting condition was retained as a separate “rest” class. In the second classification analysis, all seven tasks were classified, following the same preprocessing and classifier configuration as in the first classification analysis.

### 2.5. Statistical analysis

When data samples were of sufficient size (N>5), data distributions were first assessed for normality using the Shapiro-Wilk test and for homogeneity of variance using Levene’s test. For comparisons between groups where the normality assumption was not met, the non-parametric Mann–Whitney U test was applied. Associations between variables, such as PEC values and network class specificity, were quantified using Spearman’s rank correlation coefficient. Where applicable, results were reported with the corresponding statistical significance.

For classification-based analyses, the receiver-operating-characteristic (ROC) and precision-recall (PR) curves were tracked per class, and confusion matrices were accumulated across folds. Performance metrics (accuracy, F1-score, precision, recall, ROC-AUC, and PR-AUC) were computed both at the class level and overall weighted average. Feature importance was estimated from the absolute values of the classifier’s learned coefficients, averaged across folds. Mean normalized confusion matrix aggregated across all subjects assessed classification performance at the group level. Each subject’s confusion matrix, reflecting predicted versus true class labels, was first row-normalized, such that the entries in each row (corresponding to a true class) were divided by the total number of samples in that class. This normalization step ensures that the matrix entries represent classification proportions rather than absolute counts, thus accounting for class imbalance within each subject. The normalized confusion matrices were then summed across all subjects and divided by the total sample count per class, to produce the mean normalized confusion matrix.

## 3. Results

### 3.1. MultiPEC on networks encoding binary streams

We simulated sets of bit streams, in which we controlled the binary code of two categories of information: letters and digits. Five-bit code for each character was distributed across a set of five nodes, such that node 1 always contained the first bit of the code, node 2, the second bit, etc., consistently (Fig 3A). This way, a network contained a character’s full data model (binary code). The complete bit stream consisted of repeats of random combinations of category characters. The sets of bit streams simulated distributed encoding of information across a network of nodes. We experimented with two networks (letters and digits) and three networks (letters, digits and alphanumeric combinations), and with the 5-bit and the 6-bit code (see Methods: 2.2. Simulated binary series for controlled validation).

**Figure 3.**
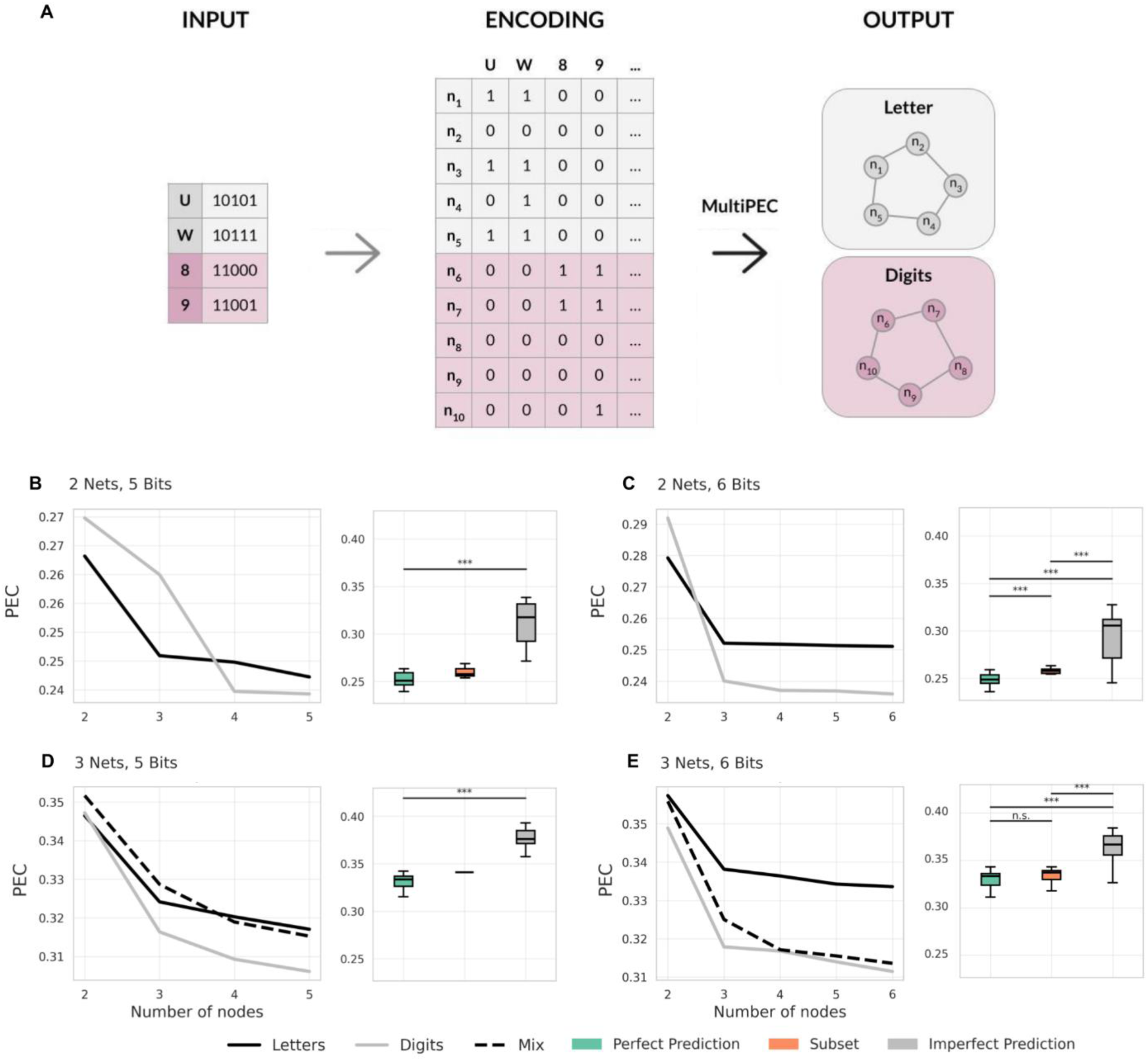
(A) Network simulation of letters and digits encoded across distributed nodes. A character was represented by a 5-bit code across 5 nodes. MultiPEC was expected to identify the letter and digit network. (B) Network PEC for two-network configuration of 5 nodes (bits), and (C) 6 bits. (D) Network PEC for three-network configuration of 5 bits, and (E) 6 bits. The left panel shows the decreasing PEC during network build-up by MultiPEC. MultiPEC processes combinations of nodes from a definite node pool, starting from node pairs and extending while the PEC value of the network is decreasing. When PEC increases, the network build-up terminates. The accurately identified networks for letters (black line), digits (gray line), and alphanumeric characters (“mix”, dashed line) are shown. The right panel shows that the accurate networks presented the lowest PEC value across configurations (perfect match; green). The identified networks were either a perfect match to the true letter/digit/mix-networks (green), a subset of the true network nodes (orange), or an imperfect match (containing nodes from more than one true network; gray). Central line indicates median score, boxes the interquartile range (IQR) from 25th to 75th percentile, whiskers 1.5 × IQR. Statistical comparisons were performed using the pairwise Mann–Whitney U test. Significance levels are indicated by asterisks: \*\*\**P* < 0.0001, “n.s.” for *P* > 0.05.

The digit network encoded 10 characters (0–9), whereas the letter network encoded 27 (the English alphabet), making the letter stream inherently less predictable. PEC values followed this trend, with the mix network, containing equal proportions of digits and letters, occupying an intermediate position (Fig. 3B–E). MultiPEC reliably recovered the data models of predefined categories (letters or digits) but also detected structure at finer and broader scales. Accurate networks were those whose identified node sets exactly matched the true letter, digit, or mix networks (Fig 3B-E, green). Subset networks occurred when MultiPEC recovered only a subset of the true nodes, corresponding to bits that were especially predictable within a code (Fig 3B-E, orange). Imperfect networks were those containing nodes drawn from more than one category (Fig 3B-E, gray), reflecting partial overlap or ambiguous predictability. Moreover, accurate networks consistently exhibited the lowest PEC values, validating PEC as a reliable biomarker of networks as information clusters.

### 3.2. MultiPEC on CNN model activations

To investigate the function of representations, we applied MultiPEC to a convolutional neural network (CNN) model, where causal manipulation is possible. We trained a CNN model on the MNIST dataset of handwritten digits (0–9; Fig 1) (Koehler, 2020). After 10 training epochs, the model achieved 98% overall test accuracy, with the lowest class accuracy of 97% (Fig S1). Kernel activations were analyzed by MultiPEC to identify MNIST data models. Network function was investigated by pruning the network nodes and analyzing the change in the pruned CNN model performance.

#### 3.2.1. Network function and class specificity

The network PEC value was strongly correlated to the PEC value of the starting node pair (Pearson r = 0.9; Fig S2). A total of 85 networks were identified. Pruned network size positively correlated with the model impairment extent (Spearman r = 0.52), but the scatterplot reveals substantial deviation from a monotonic trend. Several small networks caused disproportionately large performance drops, while some large networks with minimal impact (Fig 4A). These extreme cases indicate that the influence of a network on model performance is not solely determined by size, suggesting the functional importance of specific networks classification. Network function was quantified via class specificity, defined as the difference between maximum absolute change in class-wise accuracy and the average absolute change across all classes (Equation 1). Class specificity showed a weak but statistically significant negative correlation with PEC (Spearman r = -0.3; Fig 4B), suggesting that networks with lower PEC values tend to encode more class-specific information. In other words, low PEC marks networks that carry information critical for distinguishing between classes, whereas high PEC networks may represent more distributed or less discriminative features. To further validate this interpretation, we compared the class specificity of identified networks against 1,000 iterations of random pruning. The mean of the kernel density estimate (KDE) was significantly higher (*P* = 0.002) for MultiPEC networks, confirming that they were more class-specific than expected by chance (Fig 4C). Notably, classes with inherently low baseline performance also showed low class specificity and fewer identified networks in both MultiPEC and random pruning (Fig S3, Table S1), suggesting that the detectability of class-specific networks is constrained by the model’s underlying representational structure.

**Figure 4.**
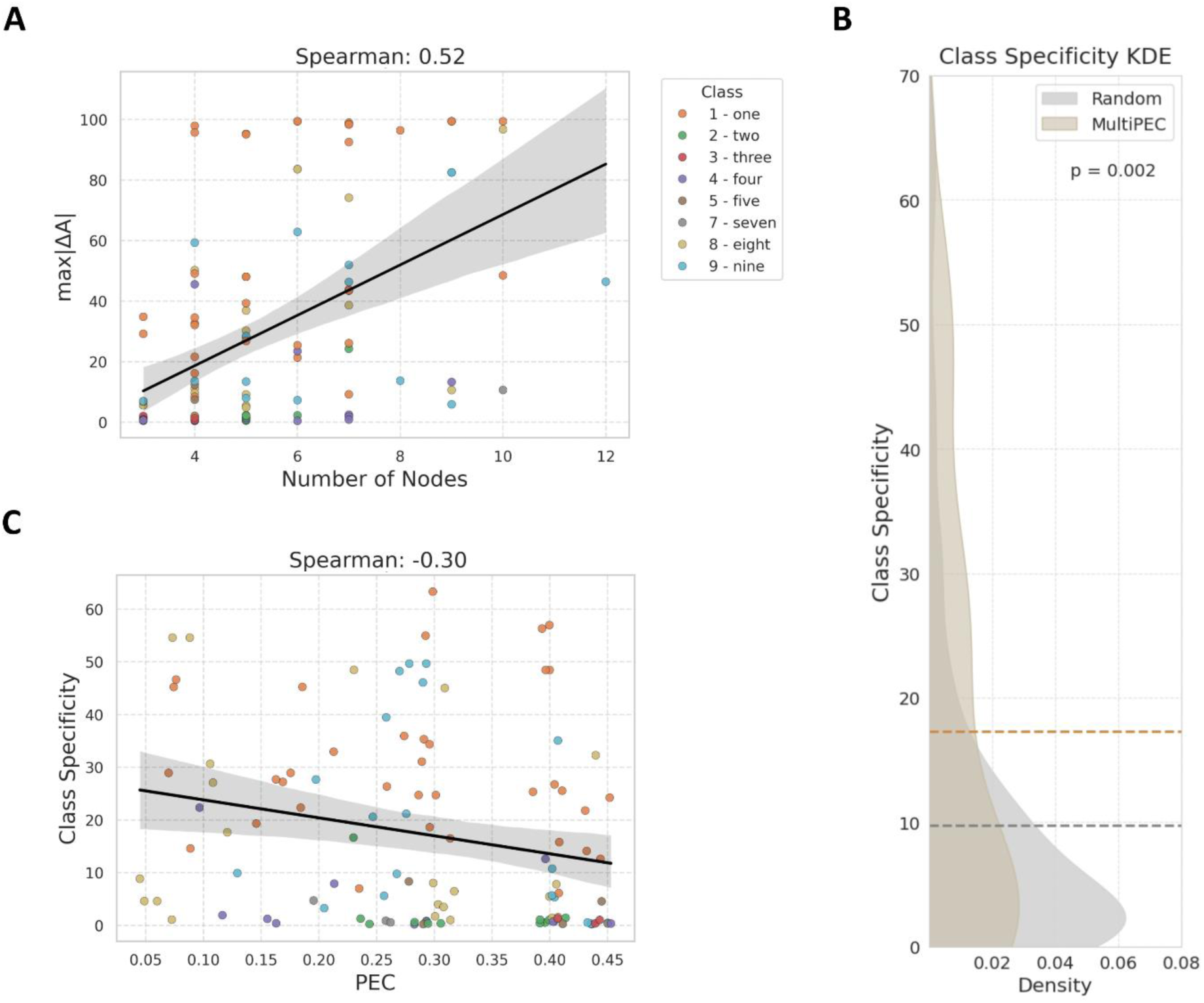
Class specificity across identified networks. (A) Relationship between network size and maximum absolute accuracy change (max|ΔA|) after network pruning. Scatter points represent individual networks, color-coded by MNIST class for which they were specific. The black regression line shows positive association (Spearman r = 0.52, *P* < 0.001), with many outliers. (B) Kernel density estimate (KDE) of class specificity distributions comparing random pruning (gray) and MultiPEC network pruning (beige). The shaded areas represent the density of specificity values across all classes. Dashed horizontal lines indicate mean specificity. Statistical significance between the two distributions was assessed using the Mann-Whitney U test (*P* = 0.002). (C) Relationship between PEC and class specificity across networks. Scatter points are represented as in (A). The black regression line illustrates a weak but significant negative linear trend (Spearman r = -0.3, *P* = 0.001).

#### 3.2.2. Data models represented by the networks

We analyzed the activation maps of low-PEC class ‘8 - eight’-specific networks (Fig 5). Pruning network {N9, N3, N5} caused a 6% drop in accuracy for class ‘8 - eight’ and 2% for class ‘9 - nine’, suggesting that the network represents features of ‘8 - eight’, which partially overlap only with class ‘9 - nine’ (Fig 5A). The activation maps show that node N9 activates strongly on upward concave contours, N3 emphasizes outer boundaries between black strokes and white background, and N5 highlights the outer edge of the first arch of a dome-shaped curve (Fig 5A). Pruning an additional node N2 reduced accuracy double for ‘8 - eight’ and ‘9 - nine’ (Fig 5B). Adding N4, which activated to internal stroke regions within curved diagonal segments, lead to a ∼40% drop in accuracy only for class ‘8 - eight’ and ∼20% for class ‘9 - nine’ (Fig 5C). Finally, adding N6 resulted in a substantial 82% drop for class ‘8 - eight’ as well as major losses in class ‘6 - six’ (59%), class ‘4 - four’ (39%), and class ‘1 - one’ (25%). Although highly specific for class ‘8 - eight’, N6 expands the network’s representational scope to features shared by several classes (outer stroke of downward concave curve; Fig 5D). This network decomposition shows that individual nodes encode geometrical features tracing a silhouette of the digit eight. MultiPEC identified node combinations that integrate data models of the input category and subcategories at varying scales. Notably, adding nodes can either reinforce class structures or dilute them by incorporating features that appear across multiple classes.

**Figure 5.**
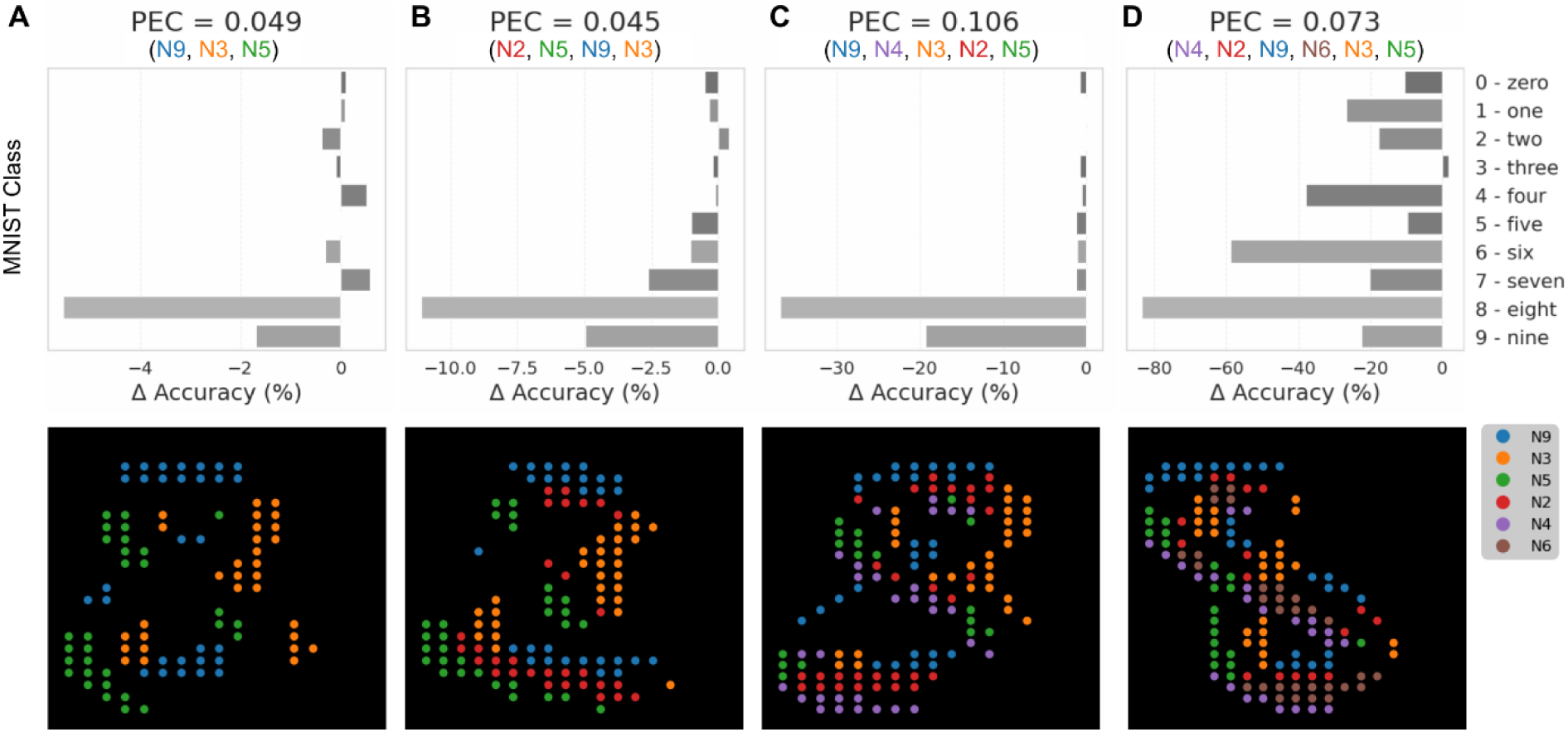
Networks specific for class ‘8 - eight’. (A) Class accuracy changes in pruned CNN models. Four selected networks showing specificity for class ‘8 - eight’ are shown from left to right in the order of increasing node count. Each successive network includes all nodes from the previous one, with additional nodes added incrementally, illustrating the data model’s complexity change with each added node. (B) Activation maps of nodes that constitute the corresponding networks from (A), in response to MNIST image of class ‘8 - eight’.

### 3.3. MultiPEC on human EEG data

We evaluated MultiPEC ability to differentiate brain networks associated with distinct perceptual conditions using publicly available EEG data. The dataset, sourced from the Open Science Framework (https://osf.io/e93db/), consisted of 64-channel EEG recordings from 24 participants who underwent seven experimental conditions. These conditions were designed to manipulate the degree of perceptual coherence and included: (1) listening to an audiobook (A1), (2) listening to a scrambled version of the audiobook (A2), (3) listening to a randomized version of the audiobook (A3), (4) watching a silent video (V1), (5) watching a scrambled version of the video (V2), (6) watching a randomized video (V3), and (7) resting with eyes open (R). It is important to note that no eye-tracking data was collected, and participants were instructed to keep their eyes open throughout the sessions. Each recording was independently processed, extracting task-specific networks for each subject.

#### 3.3.1. Classification of auditory and visual modalities in identified networks

Networks were transformed into vectors, which contained cognitive functions associated with EEG channels (Fig 2), weighted by the network’s PEC. Figure 6 illustrates the network vectors for two subjects. In Subject 1, the identified networks show a clear differentiation between the conditions (Fig 6A). The three audio conditions present activations in auditory processing. Language- and speech-related functions are active in A1 but absent in A2 and A3. However, auditory associations and sound localization remain active in A2 and A3, suggesting preserved low-level auditory processing in the absence of semantic content. In contrast, the visual conditions in the same subject present activation of object recognition and higher-order visual processing functions. These functions are most prominent in V1 condition and show a gradual reduction in the V2 and V3, where visual coherence is progressively degraded. Notably, the resting state (R) presents low activation across functions. This subject represents a case where the identified functional networks align well with the expected perceptual demands of each task, suggesting that MultiPEC captured condition-specific cognitive processes, even in coarse recordings such as scalp EEG. Despite a clear distinction of networks across conditions, the number of network samples was not sufficient for classification. Classification was feasible in 22 out of 24 subjects (e.g., Fig 6B), who had at least five networks per condition (see Methods: 2.4.3. Task classification). Some subjects presented activation of both auditory and visual functions across all conditions, which may be due to the design of the experiment, having the eyes opened during the audio conditions.

**Figure 6.**
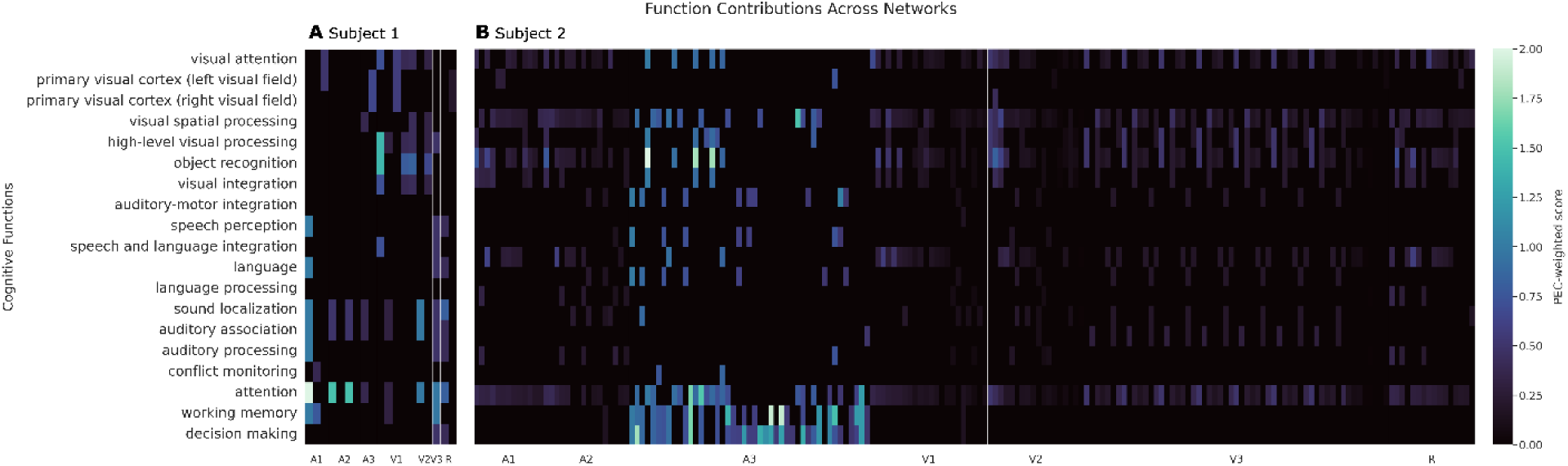
Network vectors of PEC-weighted scores for cognitive functions for (A) Subject 1 and (B) Subject 2. Rows correspond to cognitive functions, and columns to networks identified for each experimental condition. Networks are grouped by condition: A1 = original audio, A2 = scrambled audio, A3 = randomized audio, V1 = original video, V2 = scrambled video, V3 = randomized video, and R = resting state. The intensity of each cell reflects the strength of a cognitive function.

To explore whether the identified networks carried discriminative signatures of the perceptual modality, we grouped the conditions into three broader classes: Audio (A1-3), Video (V1-3), and Rest. The classification analysis was then performed using a SVM classifier to assess whether networks could reliably distinguish between these three groups. Classification performance varied across subjects (Fig 7A; precision from 0.27 to 0.87, recall from 0.23 to 0.81, F1-score from 0.25 to 0.83, ROC-AUC from 0.5 to 0.9, and PR-AUC from 0.39 to 0.87). The classifier correctly identified the Audio class in 56% of the cases, Video in 58%, and Rest in 60%, suggesting a moderate but consistent separability between these brain network profiles (Fig 7B). These results indicate that MultiPEC captured modality-specific data models, even when the underlying stimuli are semantically degraded and visual stimulus is maintained.

**Figure 7.**
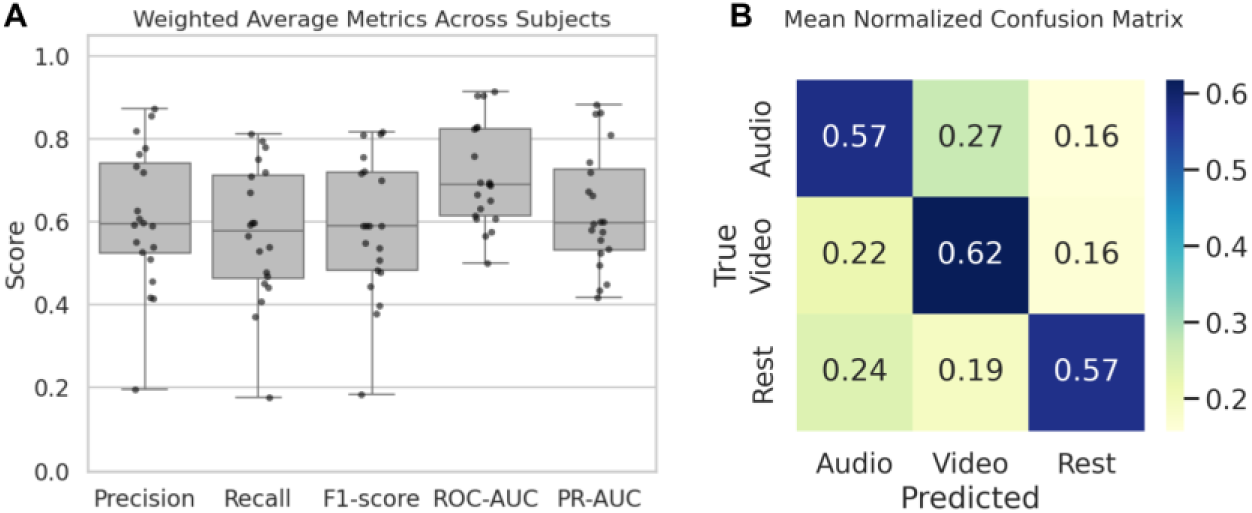
Classification performance and feature importance for the 3-class case. (A) Weighted average performance metrics (precision, recall, F1-score, ROC-AUC, and PR-AUC). Scattered points show individual subjects. Central line indicates median score, boxes indicate the IQR, whiskers indicate 1.5 × IQR. (B) Mean normalized confusion matrix across subjects.

#### 3.3.2. Classification of perceptual sub-modalities in identified networks

We next tested whether the networks could be classified into specific experimental conditions. The randomized audiobook condition (A3) achieved the highest classification scores (Fig 8A). Both randomized conditions (A3, V3) outperformed their scrambled counterparts (A2, V2). F1-scores varied across conditions (A1: 0.06 – 0.73; A2: 0.05 – 0.84; A3: 0.14 – 0.96; V1: 0 – 0.80; V2: 0.00 – 0.40; V3: 0.09 – 0.75; R: 0.00 – 0.71). Weighted average metrics revealed wide variability across subjects, with precision ranging from 0.05 to 0.84 and recall from 0.07 to 0.8 (Fig 8B). ROC-AUC values reached as high as 0.94, indicating strong discriminative ranking ability even in cases where F1-score was low. Confusion matrices confirmed that A3 was most accurately classified (68%; Fig 8C). Accuracy overcame random performance for all classes. Feature importance analysis highlighted auditory-related functions (sound localization, auditory processing, language; Fig 8D). Interestingly, non-specific functions such as motor planning and attention were also important. Visual processing functions ranked lower, consistent with the design of the experiment.

**Figure 8.**
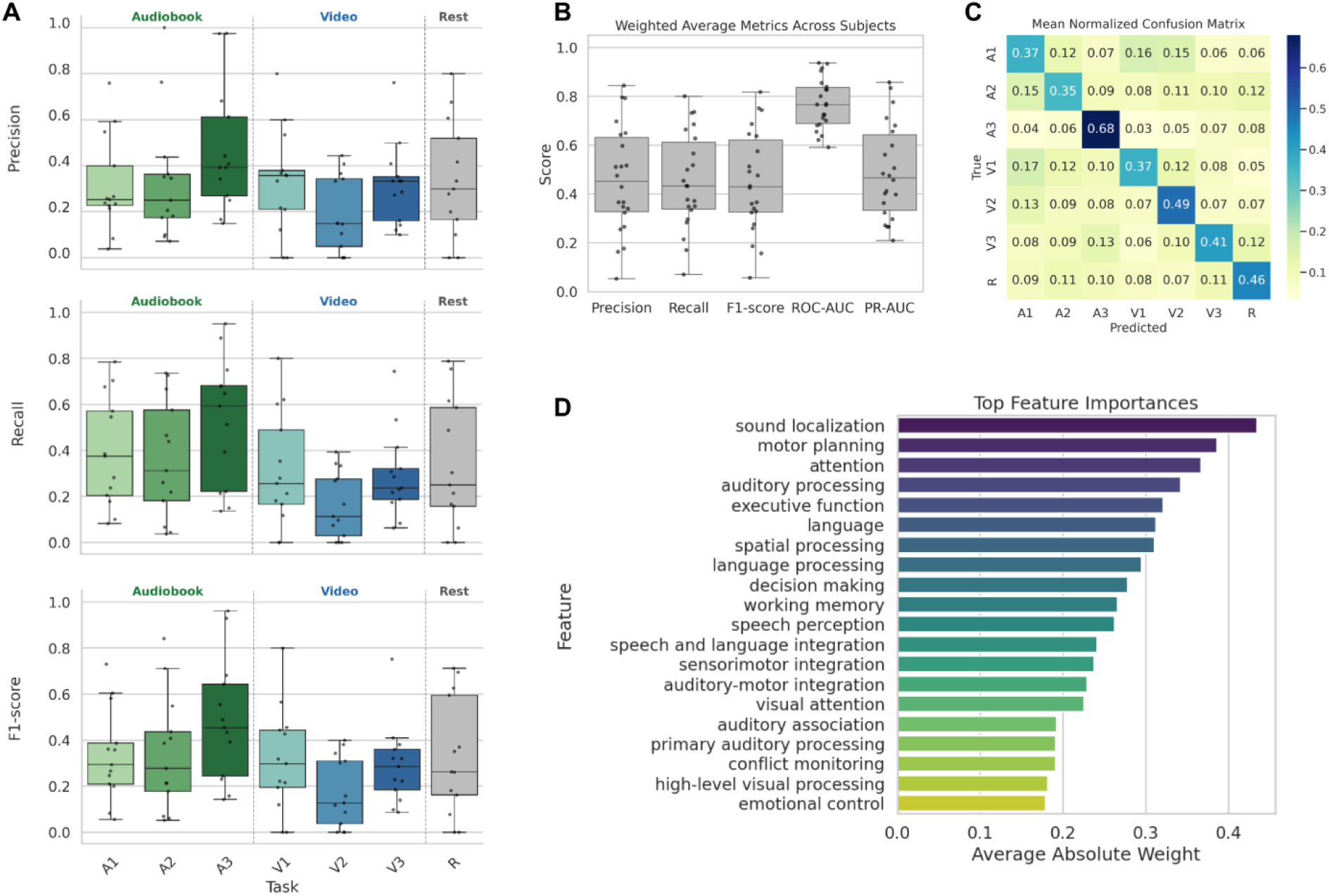
Classification performance and feature importance. (A) Precision, recall, and F1-scores for each condition. Scattered points show individual subjects, central line the median, boxes the IQR, whiskers extend 1.5 × IQR. Audiobook conditions (A1-3) are in shades of green, video conditions (V1-3) in shades of blue, and rest (R) in gray. (B) Weighted average performance metrics (precision, recall, F1-score, ROC-AUC, PR-AUC) across subjects. (C) Mean normalized confusion matrix. Classes with insufficient data were excluded. (D) Top features ranked by average absolute weight. A1 = original audio, A2 = scrambled audio, A3 = randomized audio, V1 = original video, V2 = scrambled video, V3 = randomized video, and R = resting state.

## 4. Discussion

In this study, we introduce MultiPEC, a tool for automatic detection of internal representations from neural activity data. MultiPEC leverages PEC as a network marker, which relates to the complexity of information contained in the network and its consistency across repetitions (Fig 3B) (Principe et al., 2019). The letter/digit network experiment served as a controlled proof-of-concept for demonstrating the foundations, performance, and limitations of MultiPEC. In this synthetic setting, the ground-truth activations of each character were known, enabling a direct validation of the method. MultiPEC successfully identified the correct networks corresponding to individual characters but also detected additional non-putative networks. Some of these consisted of subsets of true networks, whereas others combined elements of multiple networks. These results illustrate that MultiPEC not only recovers known information structures but also detects emergent dependencies present in the data. Importantly, this experiment revealed that the PEC value acts as a quantitative signature of information representation: true information patterns tend to exhibit lower PEC values, reflecting their stronger internal consistency and predictive structure. Understanding how to interpret the PEC values in this controlled scenario provided the foundation for extending MultiPEC to more complex and less constrained systems.

We next applied it to a CNN, a model system of parallel distributed processing that allows direct manipulation and validation of network components (Bowers, 2017). CNNs are well suited for visual tasks, particularly interesting because they allow reconstructing the learnt image patterns from node activations, which could then be used as a reference for validating MultiPEC networks (Itoh et al., 1990). CNN mimics feature extraction in the visual cortex, using kernels, learnable matrices that slide over the input data and extract local features from a local input area, like a receptive field (LeCun et al., 2010; Lindsay, 2021). Kernels consist of a population of neurons, and their activations present an artificial analogue to population-level neural responses recorded by EEG, enabling MultiPEC to be applied seamlessly both to electrophysiological time series and CNN activation data. Model pruning allowed interrogating the function of specific networks through change in model performance. Importantly, the extent of performance impairment was not solely determined by network size, as small networks caused disproportionately large drops in accuracy (Fig 4A). The identified networks were indeed meaningful functional units, responsible for specific aspects of model performance. Further supporting this, class specificity analysis showed that these networks had significantly better class specificity compared to 1,000 random pruning instances (Fig 4C), confirming that the networks represented class-relevant information beyond chance. Additionally, classes with inherently low specificity baseline also exhibited low specificity and fewer networks than in the MultiPEC results, suggesting that reduced specificity in these cases reflects the model’s underlying functional architecture rather than a methodological limitation (Fig S3, Table S1). A weak but significant negative correlation between PEC and class specificity indicates that low-PEC networks tend to represent class features, such as combinations of edges and curves composing a specific digit (Fig 5). In contrast, high-PEC networks were more likely to lack class specificity (Fig 4C). This derives from the MultiPEC algorithm feature that permits all possible combinations of nodes. Therefore, the high-PEC clusters may represent super-categories or shared regularities across classes. This interpretation aligns with the findings from the binary series experiment, where low-PEC networks encoded stable categories of information (letters and digits), while higher PEC captured more variable representations at the intersection of base categories (Fig 3C).

Having validated MultiPEC in both synthetic and artificial neural systems, we next examined whether its principles extend to biological neural data. The human brain presents a far more complex and noisy network system. Nevertheless, the ability to map functional networks associated with sensory, cognitive, or pathological states using non-invasive recordings would be transformative. Using EEG recordings from a perceptual experiment, we found that MultiPEC detects networks with functional signatures specific to broad perceptual modalities (audio, video, rest) and, at finer resolution, sub-modalities within each modality. Auditory-related cognitive functions, such as sound localization, speech perception and language processing, predominated when subjects listened to the audiobook, even though they were simultaneously exposed to visual stimuli as their eyes were opened. High-level visual processing and object recognition predominated when subjects watched a soundless video. Classification of networks improved when all seven conditions were considered, indicating that the data models more likely matched the more precise categories rather than broader input modalities, like audio processing. This finding illustrates that the definition of neural representations is influenced by the definition of the situational context. Classification was possible on subject-level, but not on group-level, indicating individualized mental models (Kahn et al., 2025; Yanagisawa et al., 2025). Inter-subject variability was reflected in the performance range, with some subjects showing clear condition-specific activation patterns, while others showed more diffuse patterns. Heterogeneity of individual experience was unavoidable due to the low-control nature of the experiment and coarse neural activity recorded by the scalp EEG (Scrivener & Reader, 2022). The stimuli were not strictly constrained, and no eye-tracking or behavioral performance measures were available, making it difficult to verify whether participants consistently attended to the stimuli or processed them in the intended manner. Still, this dataset represents a good starting point for our proof-of-concept study, as it included enough subjects, whole brain coverage, consistent recordings across subjects, and experimental conditions that could be analyzed at several modality scales (Orłowski & Bola, 2023). The most precise network definition was achieved in the randomized audiobook condition, which likely required most attention and was the most unique, given the presence of an auditory stimulus and absence of semantic context. The strong ROC-AUC scores, even when precision and recall were lower, confirmed that the network classes were generally distinguishable, though not always reliably classified across all subjects. Finally, the convergence of results across EEG and CNN experiments demonstrates that MultiPEC’s principles are generalizable, capturing task-relevant representations in both biological and artificial architectures. This finding also implies a common strategy of information storage in the two systems.

A key technical limitation of the MultiPEC emerges in models with many nodes. Because the number of possible node combinations increases exponentially, computation becomes impractical. In our analyses, we observed a strong correlation between the network PEC value and the PEC value of the starting node pair (Fig S2), suggesting that the search space could be reduced without substantial loss of sensitivity. Restricting the number of node pairs may therefore offer a computationally feasible strategy for applying PEC to large-scale networks, although at the potential cost of missing rare but functionally important combinations.

Altogether, our findings highlight PEC as an informative marker for identifying functionally relevant information clusters and underscore the importance of focusing on low-PEC ranges in the search for meaningful network representations. More broadly, MultiPEC directly addresses a central gap in current approaches to neural data. While functional connectivity, dimensionality reduction, supervised decoders, and brain foundation models can reveal associations or predictive features, they do not specify which sets of nodes compose a representation. At present, no existing method provides an equivalent operation, identifying explicit node sets that constitute an information representation, making direct benchmarking to a state-of-the-art alternative impossible, as related approaches address fundamentally different questions. MultiPEC fills this gap by providing explicit node sets that encode shared information, making neural representations empirically identifiable and directly testable. In this sense, MultiPEC contributes to the broader challenge of linking distributed activity to function, bridging the gap between statistical descriptions of neural data and mechanistic accounts of representation. Looking forward, applying MultiPEC to higher-density and multimodal recordings may help uncover how representations reorganize during learning, recovery, or neurofeedback, and how they can be leveraged for adaptive brain–computer interfaces.

## 5. Conclusion

Taken together, these findings position MultiPEC as a promising framework for mapping functional networks in both biological and artificial systems. By extracting node sets that jointly minimize prediction error, MultiPEC identifies candidate neural representations in a fully data-driven manner. This addresses a central limitation of existing methods, which can capture associations or predictive features but leave the structure of representations implicit. In this way, MultiPEC contributes to bridging the gap between statistical descriptions of neural activity and mechanistic accounts of representation.

## CRediT authorship contribution statement

Karla Ivankovic: Conceptualization, Data curation, Formal analysis, Investigation, Methodology, Resources, Software, Validation, Visualization, Writing – original draft; Anastasios Dimou: Conceptualization, Writing – review & editing, Supervision; Justo Montoya-Gálvez: Visualization, Writing – review & editing, Resources; Riccardo Zucca: Writing – review & editing; Manuel Valero: Writing – review & editing, Supervision, Conceptualization; Alessandro Principe: Conceptualization, Resources, Project administration, Supervision, Writing – review & editing.

## Data availability

In this work we used a freely available dataset from the Open Science Framework (https://osf.io/e93db/). The MultiPEC package along with the data and notebooks for reproducing the results and figures can be accessed here: https://github.com/principelab/multipec-core.git

## Acknowledgements

The authors wish to thank Mara Dierssen, Zisis Batzos and Alex Doumanoglou for their invaluable contributions and generous support. We thank Paweł Orłowski and Michał Bola for publically sharing their EEG dataset. This work received funding from Fundación Tatiana – Proyectos en Neurociencia, with reference 2022-1514.

## References

Ansel, J., Yang, E., He, H., Gimelshein, N., Jain, A., Voznesensky, M., Bao, B., Bell, P., Berard, D., Burovski, E., Chauhan, G., Chourdia, A., Constable, W., Desmaison, A., Devito, Z., Ellison, E., Feng, W., Gong, J., Gschwind, M., … Chintala, S. (2024). PyTorch 2: Faster Machine Learning Through Dynamic Python Bytecode Transformation and Graph Compilation. International Conference on Architectural Support for Programming Languages and Operating Systems - ASPLOS, 2, 929–947. 10.1145/3620665.3640366

Borst, A., & Theunissen, F. E. (1999). Information theory and neural coding. Nature Neuroscience, 2(11), 947–957. 10.1038/14731;KWRD

Bowers, J. S. (2017). Parallel Distributed Processing Theory in the Age of Deep Networks. Trends in Cognitive Sciences, 21(12), 950–961. 10.1016/J.TICS.2017.09.013

Carrillo-Reid, L., Han, S., Yang, W., Akrouh, A., & Yuste, R. (2019). Controlling Visually Guided Behavior by Holographic Recalling of Cortical Ensembles. Cell, 178(2), 447–457.e5. 10.1016/J.CELL.2019.05.045/ASSET/B505D98C-AE06-4C0C-9ECF-4741218C1E4B/MAIN.ASSETS/FIGS1.JPG

Chang, J. C., Perich, M. G., Miller, L. E., Gallego, J. A., & Clopath, C. (2024). De novo motor learning creates structure in neural activity that shapes adaptation. Nature Communications, 15(1), 1–16. 10.1038/S41467-024-48008-7;SUBJMETA

Combrisson, E., Basanisi, R., Neri, M., Auzias, G., Petri, G., Marinazzo, D., Panzeri, S., & Brovelli, A. (2025). Higher-order and distributed synergistic functional interactions encode information gain in goal-directed learning. Nature Communications 2025 16:1, 16(1), 1–19. 10.1038/S41467-025-62507-1

Cunningham, J. P., & Yu, B. M. (2014). Dimensionality reduction for large-scale neural recordings. Nature Neuroscience, 17(11), 1500–1509. 10.1038/NN.3776;SUBJMETA

Ding, Y., Udompanyawit, C., Zhang, Y., & He, B. (2025). EEG-based brain-computer interface enables real-time robotic hand control at individual finger level. Nature Communications, 16(1), 1–20. 10.1038/S41467-025-61064-X;TECHMETA

Farzmahdi, A., Kohn, A., & Coen-Cagli, R. (2025). Relating natural image statistics to patterns of response covariability in macaque primary visual cortex. Nature Communications, 16(1), 1–13. 10.1038/S41467-025-62086-1;SUBJMETA

Hubel, D. H., & Wiesel, T. N. (1962). Receptive fields, binocular interaction and functional architecture in the cat’s visual cortex. The Journal of Physiology, 160(1), 106–154. 10.1113/JPHYSIOL.1962.SP006837;PAGEGROUP:STRING:PUBLICATION

Itoh, K., Zhang, W., Ichioka, Y., & Tanida, J. (1990). Parallel distributed processing model with local space-invariant interconnections and its optical architecture. Applied Optics, Vol. 29, Issue 32, Pp. 4790-4797, 29(32), 4790–4797. 10.1364/AO.29.004790

Iurilli, G., & Datta, S. R. (2017). Population Coding in an Innately Relevant Olfactory Area. Neuron, 93(5), 1180–1197.e7. 10.1016/J.NEURON.2017.02.010/ATTACHMENT/CE4BA710-56B8-4A50-9950-ED5B68C90E60/MMC2.PDF

Kahn, A. E., Szymula, K., Loman, S., Haggerty, E. B., Nyema, N., Aguirre, G. K., & Bassett, D. S. (2025). Network structure influences the strength of learned neural representations. Nature Communications, 16(1). 10.1038/S41467-024-55459-5,

Koehler, G. D. M. P., S. J. A. A. M. P. (2020, February 17). MNIST Handwritten Digit Recognition in Keras. Https://Nextjournal.Com/Gkoehler/Digit-Recognition-with-Keras.

Koessler, L., Maillard, L., Benhadid, A., Vignal, J. P., Felblinger, J., Vespignani, H., & Braun, M. (2009). Automated cortical projection of EEG sensors: Anatomical correlation via the international 10–10 system. NeuroImage, 46(1), 64–72. 10.1016/J.NEUROIMAGE.2009.02.006

Lebedev, M. A., & Nicolelis, M. A. L. (2006). Brain-machine interfaces: past, present and future. Trends in Neurosciences, 29(9), 536–546. 10.1016/j.tins.2006.07.004

LeCun, Y., Kavukcuoglu, K., & Farabet, C. (2010). Convolutional networks and applications in vision. ISCAS 2010 - 2010 IEEE International Symposium on Circuits and Systems: Nano-Bio Circuit Fabrics and Systems, 253–256. 10.1109/ISCAS.2010.5537907

Lever, J., Krzywinski, M., & Altman, N. (2017). Points of Significance: Principal component analysis. Nature Methods, 14(7), 641–642. 10.1038/NMETH.4346;SUBJMETA

Lindsay, G. W. (2021). Convolutional Neural Networks as a Model of the Visual System: Past, Present, and Future. Journal of Cognitive Neuroscience, 33(10), 2017–2031. 10.1162/JOCN_A_01544

Liu, Z. Q., Luppi, A. I., Hansen, J. Y., Tian, Y. E., Zalesky, A., Yeo, B. T. T., Fulcher, B. D., & Misic, B. (2025). Benchmarking methods for mapping functional connectivity in the brain. Nature Methods, 22(7), 1593–1602. 10.1038/S41592-025-02704-4;SUBJMETA

Mathis, M. W., Perez Rotondo, A., Chang, E. F., Tolias, A. S., & Mathis, A. (2024). Decoding the brain: From neural representations to mechanistic models. Cell, 187(21), 5814–5832. 10.1016/J.CELL.2024.08.051

Metzger, S. L., Littlejohn, K. T., Silva, A. B., Moses, D. A., Seaton, M. P., Wang, R., Dougherty, M. E., Liu, J. R., Wu, P., Berger, M. A., Zhuravleva, I., Tu-Chan, A., Ganguly, K., Anumanchipalli, G. K., & Chang, E. F. (2023). A high-performance neuroprosthesis for speech decoding and avatar control. Nature, 620(7976), 1037–1046. 10.1038/S41586-023-06443-4;TECHMETA

Orłowski, P., & Bola, M. (2023). Sensory modality defines the relation between EEG Lempel–Ziv diversity and meaningfulness of a stimulus. Scientific Reports, 13(1), 1–9. 10.1038/S41598-023-30639-3;SUBJMETA=2613,2649,378,3917,631;KWRD=COGNITIVE+NEUROSCIENCE,SENSORY+PROCESSING,VISUAL+SYSTEM

Pandarinath, C., Gilja, V., Blabe, C. H., Nuyujukian, P., Sarma, A. A., Sorice, B. L., Eskandar, E. N., Hochberg, L. R., Henderson, J. M., & Shenoy, K. V. (2015). Neural population dynamics in human motor cortex during movements in people with ALS. ELife, 4(JUNE). 10.7554/ELIFE.07436

Panzeri, S., Macke, J. H., Gross, J., & Kayser, C. (2015). Neural population coding: combining insights from microscopic and mass signals. Trends in Cognitive Sciences, 19(3), 162–172. 10.1016/J.TICS.2015.01.002

Panzeri, S., Moroni, M., Safaai, H., & Harvey, C. D. (2022). The structures and functions of correlations in neural population codes. Nature Reviews Neuroscience, 23(9), 551–567. 10.1038/S41583-022-00606-4;SUBJMETA

Principe, A., Ley, M., Conesa, G., & Rocamora, R. (2019). Prediction error connectivity: A new method for EEG state analysis. NeuroImage. 10.1016/j.neuroimage.2018.11.052

Sadtler, P. T., Quick, K. M., Golub, M. D., Chase, S. M., Ryu, S. I., Tyler-Kabara, E. C., Yu, B. M., & Batista, A. P. (2014). Neural constraints on learning. Nature, 512(7515), 423–426. 10.1038/NATURE13665;SUBJMETA

Santoro, A., Battiston, F., Lucas, M., Petri, G., & Amico, E. (2024). Higher-order connectomics of human brain function reveals local topological signatures of task decoding, individual identification, and behavior. Nature Communications, 15(1), 1–12. 10.1038/S41467-024-54472-Y;SUBJMETA

Schreiber, K., & Krekelberg, B. (2013). The Statistical Analysis of Multi-Voxel Patterns in Functional Imaging. PLOS ONE, 8(7), e69328. 10.1371/JOURNAL.PONE.0069328

Scrivener, C. L., & Reader, A. T. (2022). Variability of EEG electrode positions and their underlying brain regions: visualizing gel artifacts from a simultaneous EEG-fMRI dataset. Brain and Behavior, 12(2). 10.1002/BRB3.2476,

Sparacino, L., Faes, L., Mijatović, G., Parla, G., Lo Re, V., Miraglia, R., de Ville de Goyet, J., & Sparacia, G. (2023). Statistical Approaches to Identify Pairwise and High-Order Brain Functional Connectivity Signatures on a Single-Subject Basis. Life, 13(10), 2075. 10.3390/LIFE13102075

Sussillo, D., Nuyujukian, P., Fan, J. M., Kao, J. C., Stavisky, S. D., Ryu, S., & Shenoy, K. (2012). A recurrent neural network for closed-loop intracortical brain–machine interface decoders. Journal of Neural Engineering, 9(2), 026027. 10.1088/1741-2560/9/2/026027

Wang, E. Y., Fahey, P. G., Ding, Z., Papadopoulos, S., Ponder, K., Weis, M. A., Chang, A., Muhammad, T., Patel, S., Ding, Z., Tran, D., Fu, J., Schneider-Mizell, C. M., da Costa, N. M., Reid, R. C., Collman, F., da Costa, N. M., Franke, K., Ecker, A. S., … Tolias, A. S. (2025). Foundation model of neural activity predicts response to new stimulus types. Nature, 640(8058), 470–477. 10.1038/S41586-025-08829-Y;TECHMETA

Yanagisawa, K., Nakai, R., Asano, K., Kashima, E. S., Sugiura, H., & Abe, N. (2025). Optimistic people are all alike: Shared neural representations supporting episodic future thinking among optimistic individuals. Proceedings of the National Academy of Sciences of the United States of America, 122(30). 10.1073/PNAS.2511101122

