## Supplementary Material for "Data-driven identification of functional networks in artificial and biological neural networks"

**Affiliations:**

Supplementary methods

Convolutional neural network (CNN) model architecture and training

The MNIST dataset contains a total of 70,000 grayscale images of 28x28 pixels, divided into the training set (60,000) and the test set (10,000). The CNN model was implemented as per (Koehler, 2020). Each convolutional layer was followed by an activation function, the rectified linear unit (ReLU) and a pooling layer. A Dropout layer was applied after conv2. The feature maps were then flattened into a 1D vector and passed through two fully connected layers to classify the input image (dimensions 320x50 and 50x10, respectively). The final output was a log SoftMax activation. This architecture was designed to balance complexity with regularization to improve generalization on unseen data. The model was implemented using PyTorch machine learning library (Ansel et al., 2024) . Training data were divided into batches of 64 images. The negative log likelihood loss function was used to measure the predicted probability distribution. Stochastic gradient descent (SGD) optimization algorithm was used to update the model parameters iteratively through backpropagation to minimize the loss function prediction error (learning rate was 0.01 and momentum was 0.5). Test data were divided into batches of 1,000 images. The model achieved an accuracy of 98% after 10 training epochs (Fig S1). Accuracy for class '0', '1' and '4' was 99%, class '2', '3', '5' and '6' was 98%, and class '7', '8' and '9' were 97%.

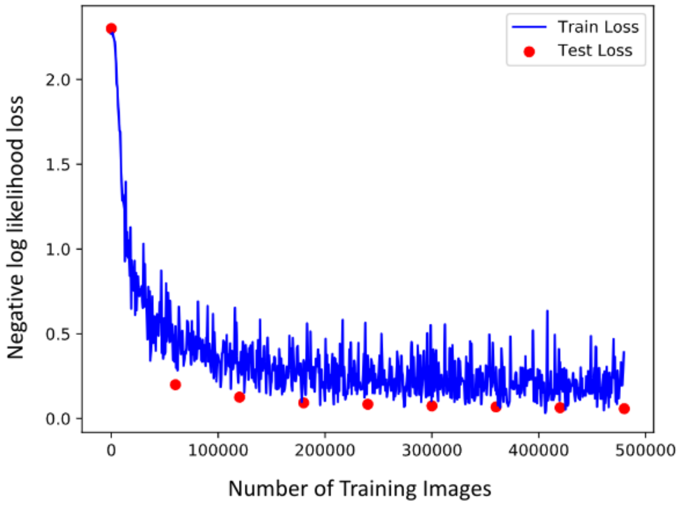


Figure S1. The prediction error calculated using the negative log likelihood loss function was continuously decreasing during training, ensuring proper training without over-fitting. Training

loss is shown after each training batch (blue line). Test loss was computed after each training epoch (red dots). The model achieved 98% accuracy on MNIST dataset after 10 training epochs.

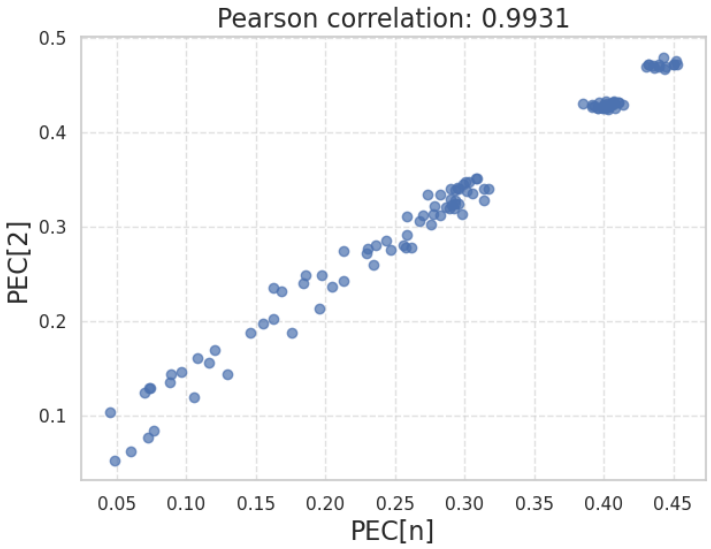


Figure S2. Relationship between pairwise PEC values (PEC[2]) and PEC values of the networks built from the node pair seed (PEC[n]), from the CNN model analysis. A strong positive linear relationship is observed, with a Pearson correlation coefficient of 0.99 ($p < 0.001$), indicating that pairwise PEC is highly predictive of the eventual network PEC.

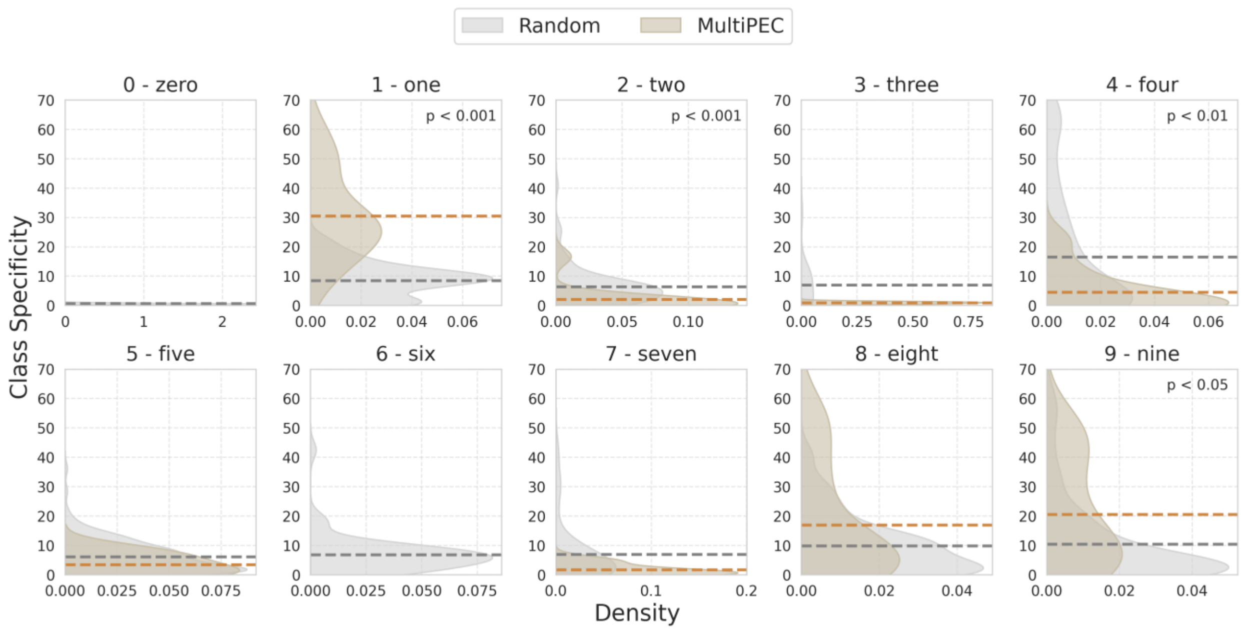


Figure S3. Per-class kernel density estimate (KDE) plots of class specificity for random pruning (gray) and MultiPEC-identified networks pruning (beige) across MNIST classes. Each subplot shows the distribution of specificity values for a single class. Dashed vertical lines mark the mean specificity for random pruning (gray) and MultiPEC network pruning (beige). Statistical significance between the two distributions was assessed per class using the Mann-Whitney U test. Significant differences are annotated in each subplot. The number of data points for each subplot are noted in Table S1.

Table S1. Number of class-specific networks identified by MultiPEC and random pruning (1,000 iterations) and maximum specificity achieved (S). MultiPEC outperforms random pruning in classes ‘1- one’ and ‘8 - eight’.

| Class | $N(Net_{Random})$ | $N(Net_{MultiPEC})$ | $S_{Random}$ | $S_{MultiPEC}$ |
|---|---|---|---|---|
| 0 - zero | 5 | 0 | 0.9 | 0.0 |
| 1 - one | 59 | 36 | 22.5 | 63.3 |
| 2 - two | 94 | 13 | 41.1 | 16.7 |
| 3 - three | 20 | 4 | 31.6 | 1.3 |
| 4 - four | 206 | 11 | 68.8 | 22.4 |
| 5 - five | 184 | 4 | 36.2 | 8.3 |
| 6 - six | 54 | 0 | 42.5 | 0.0 |
| 7 - seven | 59 | 4 | 43.8 | 4.7 |
| 8 - eight | 110 | 22 | 41.9 | 54.6 |
| 9 - nine | 209 | 19 | 64.5 | 49.7 |
